# Exploring phylogenomic relationships within Myriapoda: should high matrix occupancy be the goal?

**DOI:** 10.1101/030973

**Authors:** Rosa Fernández, Gregory D. Edgecombe, Gonzalo Giribet

## Abstract

Myriapods are one of the dominant terrestrial arthropod groups including the diverse and familiar centipedes and millipedes. Although molecular evidence has shown that Myriapoda is monophyletic, its internal phylogeny remains contentious and understudied, especially when compared to those of Chelicerata and Hexapoda. Until now, efforts have focused on taxon sampling (e.g., by including a handful of genes in many species) or on maximizing matrix occupancy (e.g., by including hundreds or thousands of genes in just a few species), but a phylogeny maximizing sampling at both levels remains elusive. In this study, we analyzed forty Illumina transcriptomes representing three myriapod classes (Diplopoda, Chilopoda and Symphyla); twenty-five transcriptomes were newly sequenced to maximize representation at the ordinal level in Diplopoda and at the family level in Chilopoda. Eight supermatrices were constructed to explore the effect of several potential phylogenetic biases (e.g., rate of evolution, heterotachy) at three levels of mean gene occupancy per taxon (50%, 75% and 90%). Analyses based on maximum likelihood and Bayesian mixture models retrieved monophyly of each myriapod class, and resulted in two alternative phylogenetic positions for Symphyla, as sister group to Diplopoda + Chilopoda, or closer to Diplopoda, the latter hypothesis having been traditionally supported by morphology. Within centipedes, all orders were well supported, but two nodes remained in conflict in the different analyses despite dense taxon sampling at the family level, situating the order Scolopendromorpha as sister group to a morphologically-anomalous grouping of Lithobiomorpha + Geophilomorpha in a subset of analyses. Interestingly, this anomalous result was obtained for all analyses conducted with the most complete matrix (90% of occupancy), being at odds not only with the sparser but more gene-rich supermatrices (75% and 50% supermatrices) or with the matrices optimizing phylogenegic informativeness and the most conserved genes, but also with previous hypotheses based on morphology, development or other molecular data sets. We discuss the implications of these findings in the context of the ever more prevalent quest for completeness in phylogenomic studies. [Chilopoda; Diplopoda; Symphyla: gene tree; species tree; node calibration; missing data.]

---

The status and interrelationships of Myriapoda have been major questions in arthropod systematics for more than a century (see a recent review in Edgecombe 2011). Variably argued to be mono-, para- or polyphyletic, molecular systematics has decisively weighed in in favor of myriapod monophyly (Regier et al. 2008; Regier et al. 2010; Rehm et al. 2014). The closest relative of Myriapoda remains a topic of discussion, though most recent analyses of arthropod phylogeny support an insect-crustacean clade named Pancrustacea or Tetraconata as sister group of Myriapoda under the Mandibulata hypothesis (Regier et al. 2010; Rota-Stabelli et al. 2011; Zwick et al. 2012; Rehm et al. 2014), a result also supported in earlier combined analyses of molecules and morphology (e.g., Giribet et al. 2001). The main alternative is an alliance between Myriapoda and Chelicerata (Paradoxopoda or Myriochelata hypotheses), retrieved in many analyses in the 1990s and 2000s. Debate has now largely shifted to the interrelationships between the four main clades of Myriapoda – Chilopoda (centipedes), Diplopoda (millipedes), Pauropoda and Symphyla – and the branching patterns of the two diverse groups, Chilopoda and Diplopoda.

The most widely endorsed phylogenetic framework for myriapods based on morphology and development divides the group into Chilopoda and a putative clade named Progoneata based on its anteriorly situated gonopore. The standard resolution within Progoneata is Symphyla as sister group of Pauropoda and Diplopoda, the latter two grouped as Dignatha (Dohle 1980, 1997; Edgecombe 2011). Progoneata has found some support from molecular phylogenetics (Gai et al. 2008; Regier et al. 2010) but it has also been contradicted by a grouping of Chilopoda and Diplopoda (Rehm et al. 2014). The most novel hypothesis to emerge from molecular data is an unanticipated union of pauropods and symphylans (Dong et al. 2012), a group that has been formalized as Edafopoda (Zwick et al. 2012).

Pauropoda and Symphyla are relatively small groups (835 and 195 species, respectively), with few taxonomic specialists. This contrasts with the more diverse and intensely studied Chilopoda and Diplopoda (Fig. 1). Of these, Chilopoda is the more extensively analyzed phylogenetically, the relationships within its four diverse orders having been evaluated based on morphology and targeted sequencing of a few markers (e.g., Giribet and Edgecombe 2013; Vahtera et al. 2013; Bonato et al. 2014a). The interrelationships of the five extant centipede orders had converged on a morphological solution (Edgecombe and Giribet 2004) that differs from the first transcriptomic analysis to sample all five orders (Fernández et al. 2014b) in one key respect: morphology unites Craterostigmomorpha (an order composed of two species of the genus *Craterostigmus*) with the two orders that have strictly epimorphic development, whereas molecules assign the order Lithobiomorpha to that position. Millipedes are the most diverse group of myriapods, with 7753 (Shear 2011) to more than 12,000 (Brewer et al. 2012) named species classified in ca. 16 orders (Brewer and Bond 2013). Millipede phylogeny has until recently been inferred based on small sets of morphological characters (Enghoff 1984; Sierwald et al. 2003; Blanke and Wesener 2014), but some analyses of a few nuclear protein-coding genes have allowed traditional ordinal-level systematics to be tested with molecular data (Regier et al. 2005; Miyazawa et al. 2014).

**Figure 1.**
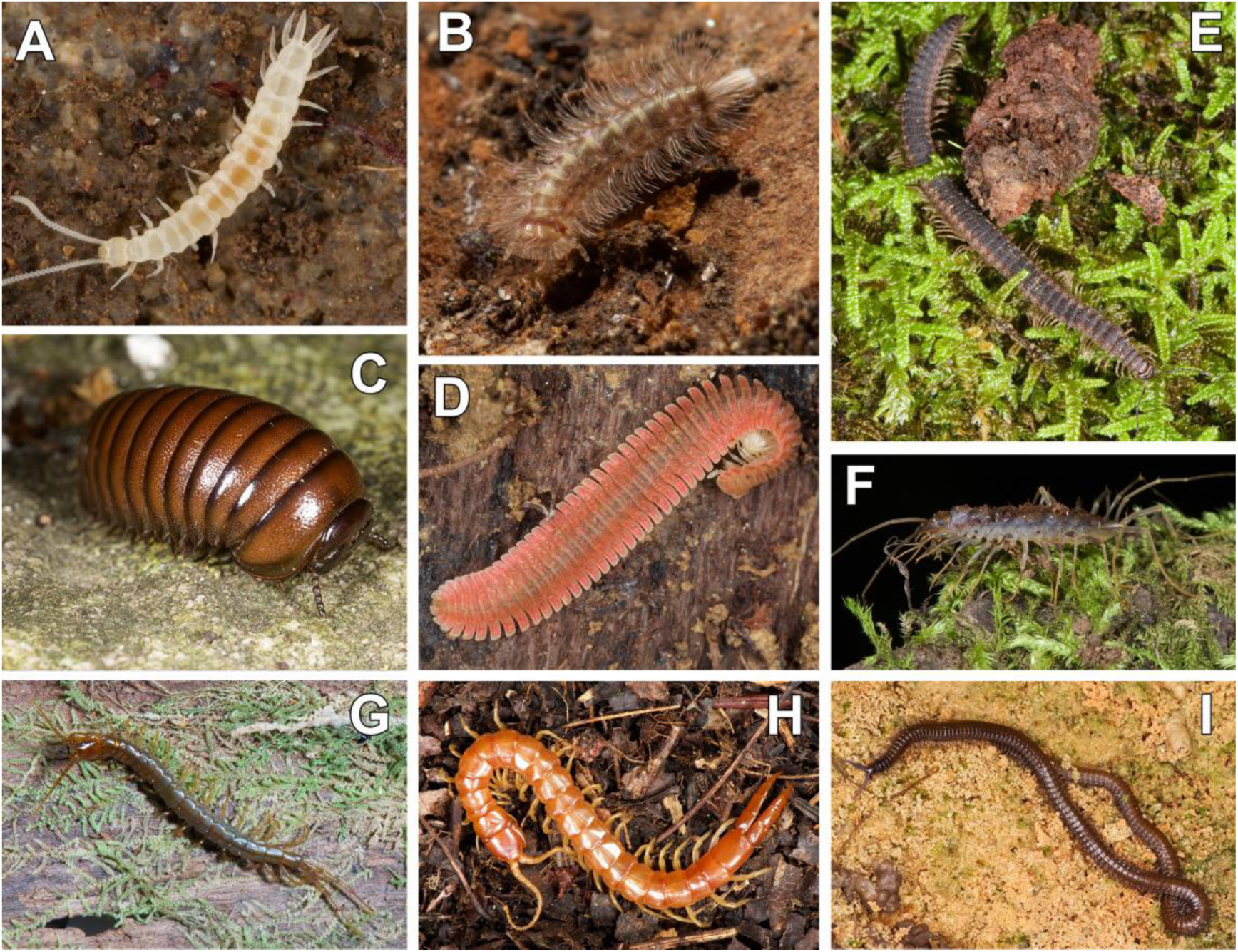
Live habitus of myriapod exemplars from this study. A, *Scutigerella* sp. from Great Smoky Mountains National Park, Tennessee, USA; B, *Eudigraphis taiwaniensis,* from Kenting National Park, Taiwan; C, Sphaerotheriid from Helderberg Nature Reserve, South Africa; D, *Brachycybe lecontii* from Great Smoky Mountains National Park, E, *Abacion* sp. from Great Smoky Mountains National Park; F, *Scutigerina weberi* from KwaZulu-Natal, South Africa; G, *Craterostigmus crabilli* from Kahurangi National Park, New Zealand; H, *Theatops spinicaudus* from Great Smoky Mountains National Park; I, *Notiphilidesgrandis* from Reserva Ducke, Manaus, Amazonas, Brazil.

Until very recently, analyses of arthropod phylogeny based on transcriptomic data have included very few myriapods, typically one species of Diplopoda and one of Chilopoda (Meusemann et al. 2010; Andrew 2011). This undersampling of myriapod diversity is now being rectified, with the first phylogenomic studies aimed at testing major hypotheses about diplopod and chilopod phylogeny appearing. Relationships between eight of the 16 recognized millipede orders have been inferred using 221 genes sampled for nine species (Brewer and Bond 2013), and interrelations of the five extant chilopod orders have been estimated using transcriptomes for seven species and up to 1,934 genes (Fernández et al. 2014b). Rehm et al. (2014) appraised the position of Myriapoda within Arthropoda using up to 181 genes from 8 myriapods, including the first transcriptomic data for Symphyla and some key millipede groups, notably Penicillata (bristly millipedes) and Glomerida (pill millipedes), the latter millipede orders not sampled by Brewer and Bond (2013). However, none of these studies included a dense taxonomic sampling across Myriapoda: to date, the maximum number of myriapods included in phylogenomic studies is ten, one centipede and nine millipedes (Brewer and Bond 2013).

Myriapods include the oldest fossil remains of terrestrial animals, with millipedes and centipedes both having records as far back as the Silurian (Wilson and Anderson 2004; Shear and Edgecombe 2010), and trace fossils indicate the activity of millipedes in the Ordovician (Wilson 2006). Silurian millipedes exhibit the earliest direct evidence for air breathing in the form of spiracles (Wilson and Anderson 2004). This geological antiquity has brought Myriapoda to the forefront in considerations of the timing of terrestrialisation in animals and plants (Kenrick et al. 2012). Such questions of timing naturally intersect with phylogenetics in the realm of molecular dating. Recent timetrees for myriapods in the context of arthropods as a whole estimate the origin of Myriapoda by the early Cambrian and the divergence of its four classes in the late Cambrian (Rota-Stabelli et al. 2011; Rehm et al. 2014), thus predicting substantial gaps in the early fossil record of the clade.

The open questions about how the main groups of myriapods are related however limit how we interpret the timing of the diversification of a major component of the soil arthropod biota. We contribute to this discussion by presenting a large injection of novel Illumina transcriptome data for myriapods, including previously unsampled millipede orders and a substantially expanded taxonomic coverage for centipedes. Furthermore, we investigate missing data in relation to potential confounding factors in phylogenomic reconstruction (e.g., evolutionary rate, compositional heterogeneity, long branch attraction and heterotachy) to dissect the individual effect of each factor on the recovered phylogenies. Concurrently, we present a morphological character set for the same set of species as sampled transcriptomically and code a set of key fossil species for their preserved morphological characters. These datasets permit diversification time to be explored for Myriapoda.

## Material and Methods

### Sample collection and molecular techniques

Twenty-five species representing three of the four major groups of myriapods (Diplopoda, Chilopoda and Symphyla) were collected and newly sequenced for this study. Our sampling was designed to maximize representation at the ordinal level in millipedes and at the family level in centipedes. Information on sampling localities can be found in Table 1, and collection details can be found online in the Museum of Comparative Zoology collections database MCZbase (http://mczbase.mcz.harvard.edu). In addition, eight millipedes (Brewer and Bond 2013), seven centipedes (six transcriptomes from Fernández et al. (2014b) and a genome from Chipman et al. (2014)) were retrieved from the Sequence Read Archive (SRA). The following taxa were used as outgroups: an onychophoran (*Peripatopsis overbergiensis*), two crustaceans (*Calanus finmarchicus* and *Daphnia pulex*), an insect (*Drosophila melanogaster*), and six chelicerates (*Limulus polyphemus, Liphistius malayanus, Damon variegatus, Mastigoproctus giganteus, Centruroides vittatus* and *Metasiro americanus*). In addition, a seventh chelicerate outgroup, *Anoplodactylus insignis* (Pycnogonida), was newly sequenced for this study. All sequenced cDNA libraries are accessioned in the Sequence Read Archive (Table 1). Tissue preservation and RNA sequencing are as described in Fernández et al. (2014b). All data sets included in this study were sequenced with the Illumina platform. In addition, we retrieved all available sequence data from the pauropod *Euripauropus spinosus* (Regier et al. 2010) in an attempt to place this group in the myriapod tree. However, only 4 genes (out of the 57 Sanger-sequenced genes) were recovered as orthologs with a minimum of 50% of gene occupancy, and the tree showed low support values (see Fig. S1).

**Table 1.**
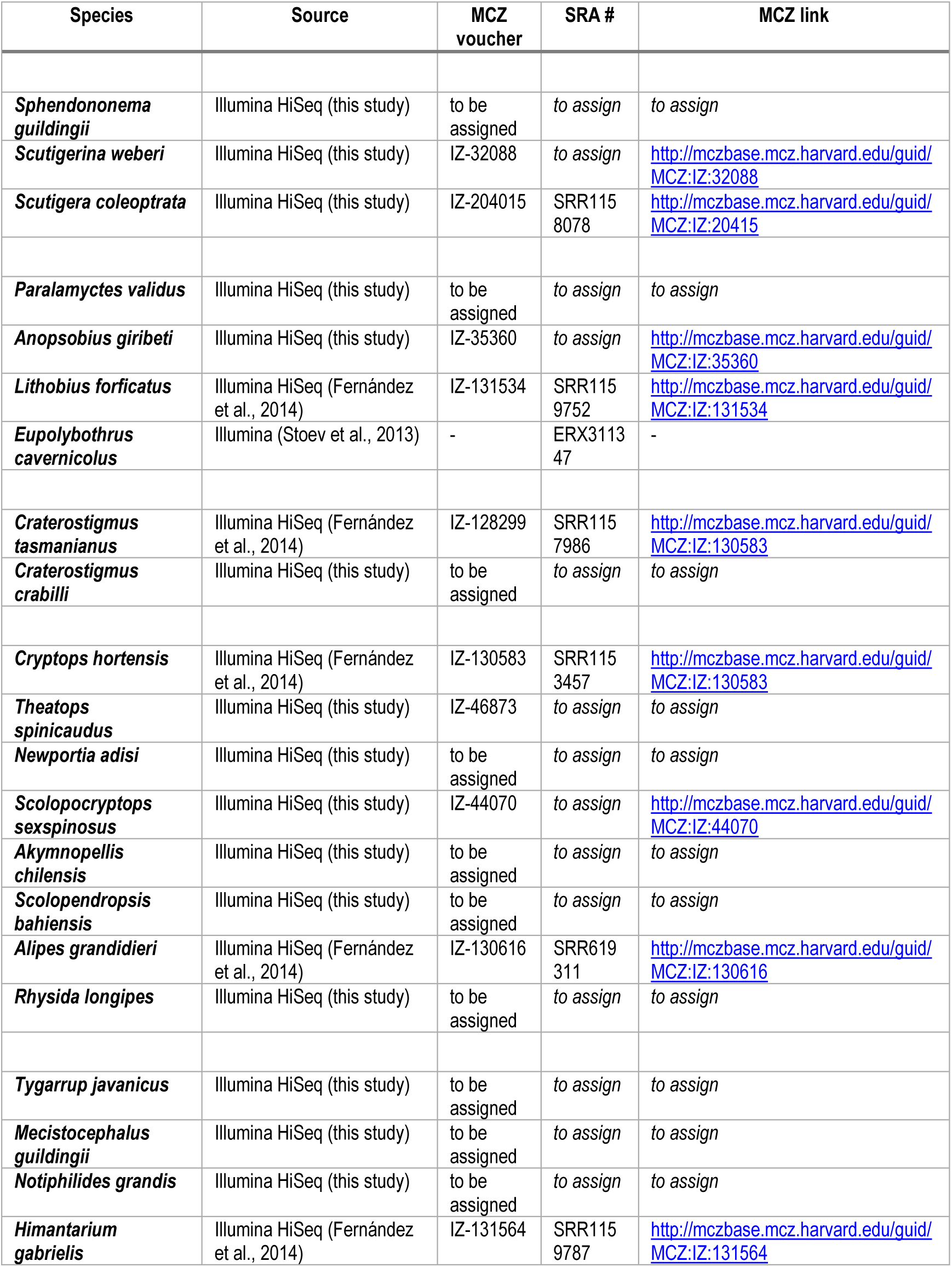
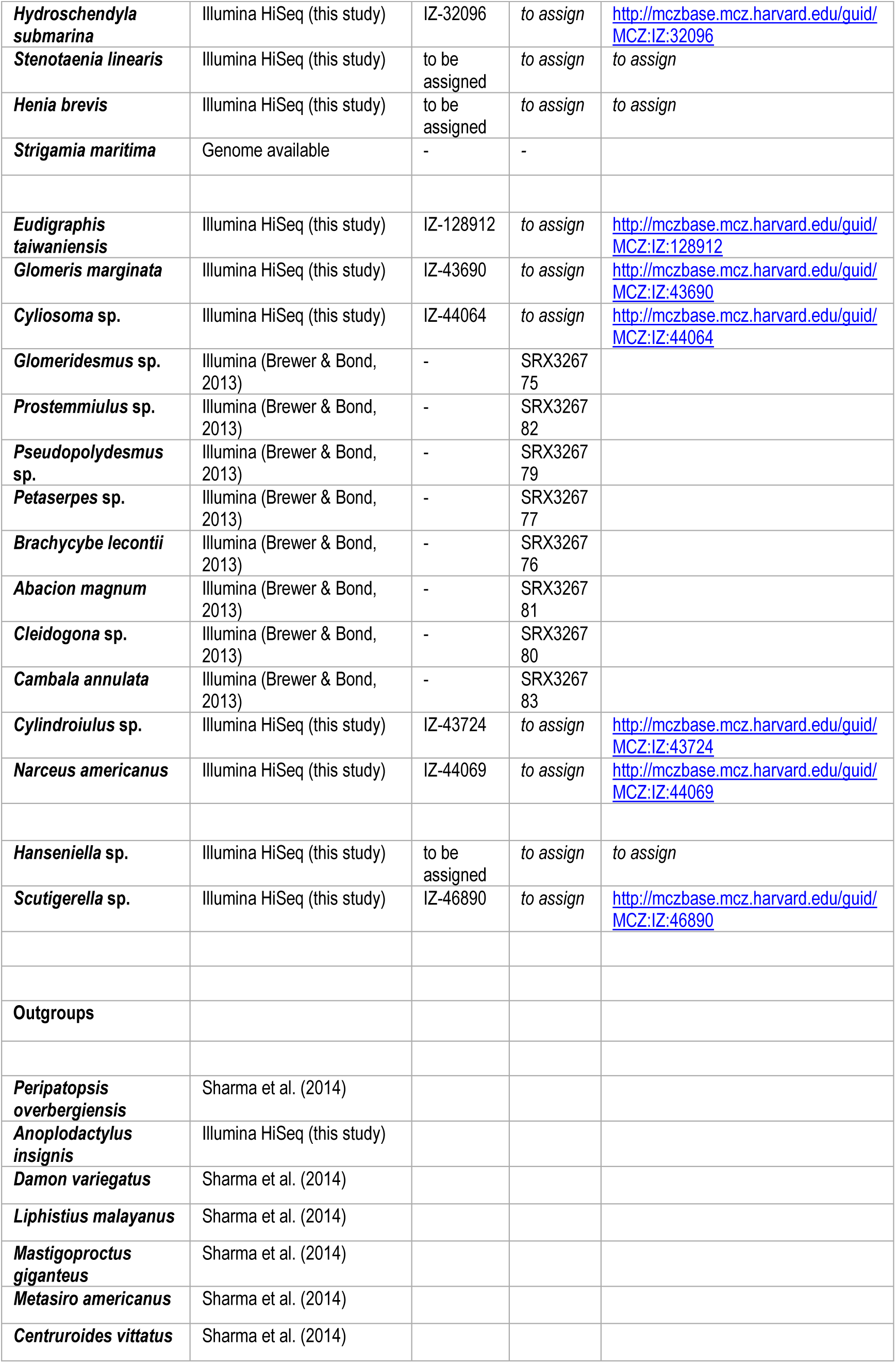
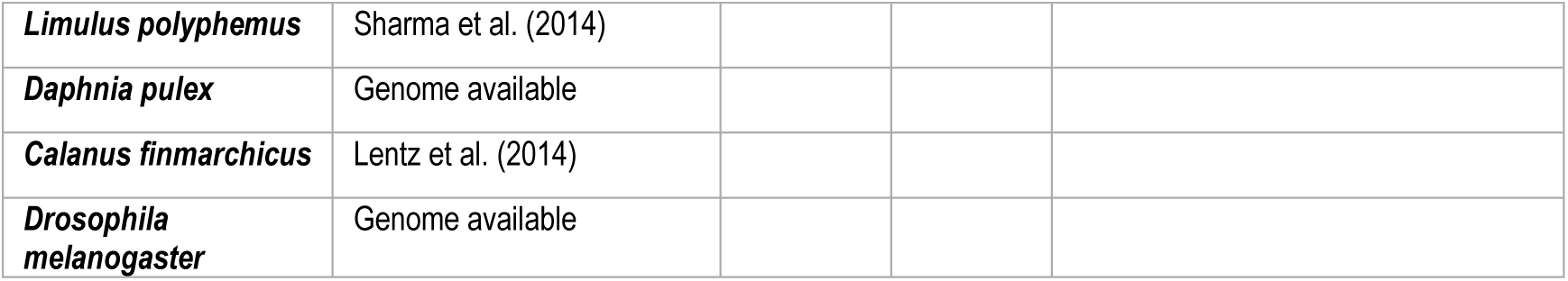
List of specimens sequenced and analyzed in the present study. The catalogue numbers and links to the Museum of Comparative Zoology (MCZ) and Sequence Read Archive (SRA) accession numbers are shown.

### Data sanitation, sequence assembly and orthology assignment

De-multiplexed Illumina HiSeq 2500 sequencing results, in FASTQ format, were retrieved from the sequencing facility via FTP. Sequenced results were quality filtered accordingly to a threshold average quality Phred score of 30 and adaptors trimmed using Trimgalore v 0.3.3 (Wu et al. 2011). Ribosomal RNA (rRNA) was filtered out. All known metazoan rRNA sequences were downloaded from GenBank and formatted into bowtie index using ‘bowtie-build’. Each sample was sequentially aligned to the index allowing up to two mismatches via Bowtie 1.0.0 (Langmead et al. 2009). Strand-specific *de novo* assemblies were done individually for each specimen in Trinity (Haas et al. 2013) with a path reinforcement distance of 75. Redundancy reduction was done with CD-HIT (Fu et al. 2012) in the raw assemblies (95% similarity). Resulting assemblies were processed in TransDecoder (Haas et al. 2013) in order to identify candidate open reading frames (ORFs) within the transcripts. Predicted peptides were then processed with a further filter to select only one peptide per putative unigene, by choosing the longest ORF per Trinity subcomponent with a python script (‘choose_longest_v3.py’, http://github//rfernandezgarcia//phylogenomics), thus removing the variation in the coding regions of our assemblies due to alternative splicing, closely-related paralogs, and allelic diversity. Peptide sequences with all final candidate ORFs were retained as multi-fasta files.

We assigned predicted ORFs into orthologous groups across all samples using the Orthologous Matrix algorithm, OMA stand-alone v0.99z (Altenhoff et al. 2011; Altenhoff et al. 2013), which has been shown to outperform alternative approaches (such as reciprocal best hit) in identifying true orthologs and minimizing Type I error in orthology assignment (Altenhoff and Dessimoz 2009).

### Phylogenomic analyses and congruence assessment

In order to explore the trade-off between number of genes and matrix completeness, three supermatrices were constructed by varying gene occupancy thresholds: supermatrix I (50% mean gene occupancy, 2,131 genes, 638,722 amino acids), supermatrix II (75% mean gene occupancy, 789 genes, 104,535 amino acids) and supermatrix III (90% gene occupancy, 123 genes, 27,217 amino acids). We also constructed two additional matrices (supermatrix IV, 87% mean gene occupancy, 40 genes, 12,348 amino acids, and supermatrix V, 87.4% mean gene occupancy, 62 genes, 16,324 amino acids) from subsets of supermatrix III, by excluding genes with the lowest phylogenetic informativeness (Townsend 2007; López-Giráldez et al. 2013). For this purpose, we analyzed the signal-to-noise distribution of the 123 genes of supermatrix III using the method described in Townsend et al. (2012) as implemented in the PhyDesign web server. This method estimates the state space and the evolutionary rates of characters to approximate the probability of phylogenetic signal versus noise due to convergence or parallelism. At every site, based on the rate of character evolution and the character state space, phylogenetic signal is characterized by the probability of observing a parsimony informative synapomorphic site pattern at the leaves of the taxa, while phylogenetic noise is characterized by the probability distribution function over time for homoplastic site patterns that mimic the correct pattern and mislead analyses. In this context, to construct supermatrices IV and V we selected the orthogroups in which the signal was higher than the noise for the shortest internodes and the longest branches in our dataset (*t_0_* = 10, *T* = 455), given an ultrametric tree generated by node calibration as described below in this section. Supermatrix IV included the 40 genes with the highest values of phylogenetic informativeness (see Fig. S2). Likewise, to construct supermatrix V we included only the genes in the two upper quartiles in their ranking of phylogenetic informativeness (62 orthologs).

Finally, to account for the effect of evolutionary rate, we ordered the orthogroups in supermatrix I based on their increasing rate and selected the 100 most conserved genes (supermatrix VI; 60.3-99.3% of conserved sites, 70% mean gene occupancy; 29,039 amino acids), the 100 genes with a variation close to the mean (supermatrix VII; 21-23.8% of conserved sites; 61% mean gene occupancy, 30,585 amino acids) and the 100 most variable genes (supermatrix VIII, 0.9-6.3% of conserved sites; 43.6% mean gene occupancy, 25,666 amino acids). Similar criteria have been applied to construct matrices for exploratory purposes in other phylogenomic analyses (e.g., Fernández et al. 2014a; Andrade et al. 2015).

The selected orthogroups in supermatrices I, II and III were aligned individually using MUSCLE version 3.6 (Edgar 2004). We then applied a probabilistic character masking with ZORRO (Wu et al. 2012) to account for alignment uncertainty, using default parameters. ZORRO assigns confidence scores ranging from 1 to 10 to each alignment site using a pair hidden Markov model (pHMM) framework. We discarded the positions assigned a confidence score below a threshold of 5 with a custom python script (‘cut_zorro.py’). These steps were not necessary in the remaining matrices as they were derived from supermatrices I and III. The aligned, masked orthogroups were then concatenated using Phyutility 2.6 (Smith and Dunn 2008).

Maximum likelihood inference was conducted with PhyML-PCMA (Zoller and Schneider 2013) with 20 nodes, and with ExaML (Aberer and Stamatakis 2013) using the per-site rate (PSR) category model. In order to further test for the effect of heterotachy and heterogeneous substitution rates, we also analyzed some of the matrices in PhyML v.3.0.3 implementing the integrated length (IL) approach (Guindon and Gascuel 2003; Guindon 2013). In this analysis, the starting tree was set to the optimal parsimony tree and the FreeRate model (Soubrier et al. 2012) was selected. Bayesian analyses were conducted with ExaBayes (Stamatakis 2014) and PhyloBayes MPI 1.4e (Lartillot et al. 2013), selecting in this last analysis the site-heterogeneous CAT-GTR model of amino acid substitution (Lartillot and Philippe 2004). Two independent MCMC chains were run for 5,000-10,000 cycles. The initial 25% of trees sampled in each MCMC run prior to convergence (judged when maximum bipartition discrepancies across chains were < 0.1) were discarded as the burn-in period. A 50% majority-rule consensus tree was then computed from the remaining trees. For practical reasons and due to the similar results obtained for the different phylogenetic analysis (see Results and Discussion), not all the analyses were implemented in all the supermatrices (see Fig. 2), but at least one ML and one Bayesian inference analysis per supermatrix were explored (Fig. 2).

**Figure 2.**
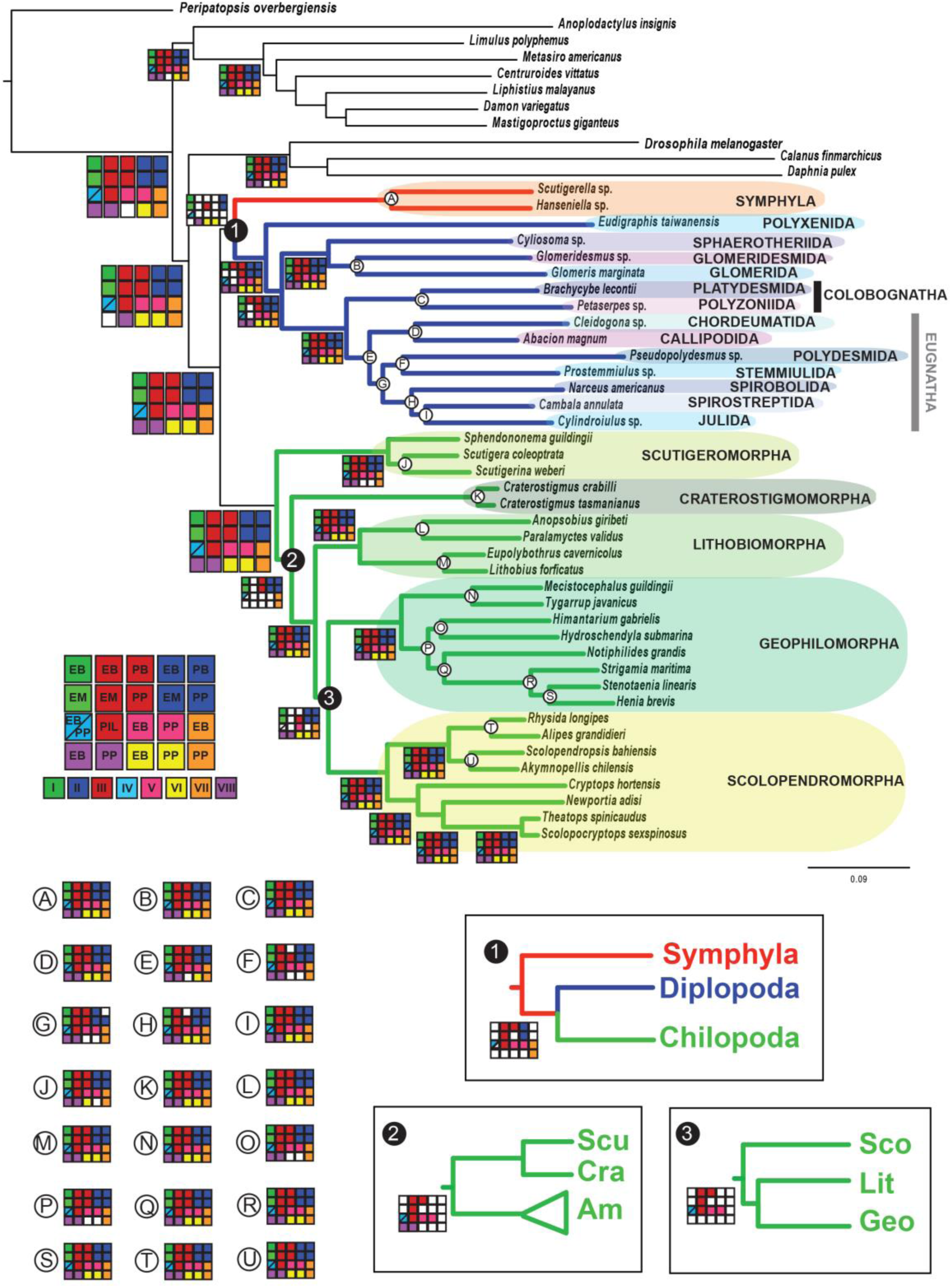
Summary of the evolutionary relationships among myriapods. Up. Maximum likelihood phylogenomic hypothesis of myriapod interrelationships (supermatrix I, Ln L ExaML-15764355.248567). Checked matrices in each node represent nodal support for the different analyses in supermatrices I to VIII (see Material and Methods for further information). PP, PhyML-PCMA; EM, ExaML; PB, PhyloBayes; EB, ExaBayes; and PIL, ML analysis with integrated branch length as implemented in PhyML. Filled squares indicate nodal support values higher than 0.95/ 0.90/95 (posterior probability, PB and EB/Shimodaira-Hasegawa-like support, PP/bootstrap, EM and PIL). White squares indicate lower nodal support. Nodal support for the clades represented with letters A to U, and the alternative topologies to the three conflicting nodes (named 1, 2 and 3) are shown.

To discern whether compositional heterogeneity among taxa and/or within each individual ortholog alignment was affecting the phylogenetic results, we further analyzed matrices I and II in BaCoCa v.1.1 (Kück and Struck 2014). The relative composition frequency variability (RCFV) measures the absolute deviation from the mean for each amino acid for each taxon. RCFV values were plotted in a heatmap using the R package gplots with an R script modified from Kück and Struck (2014).

To investigate potential incongruence between individual gene trees, we inferred gene trees for each OMA group included supermatrices I, II and III (Fernández et al. 2014a; Fernández et al. 2014b; Laumer et al. 2015). Best-scoring ML trees were inferred for each gene under the selected model of sequence evolution from 100 replicates. Gene trees were decomposed into quartets with SuperQ v.1.1 (Grünewald et al. 2013), and a supernetwork assigning edge lengths based on quartet frequencies was inferred selecting the “balanced” edge-weight optimization function, applying no filter; the supernetworks were visualized in SplitsTree v.4.13.1 (Huson and Bryant 2006). All Python custom scripts can be downloaded from https://github.com/rfernandezgarcia/phylogenomics.

As our data set includes several ancient lineages (in the order of several hundreds of Ma) with a low diversity of extant species, we further evaluated the potential effect of long branch attraction (LBA) in three lineages that show less extant diversity than their extant closest relatives: Craterostigmomorpha, Scutigeromorpha and Polyxenida. Previous morphological and molecular analyses have recovered Scutigeromorpha as the sister group to all other centipede orders (e.g., Shear and Bonamo 1988; Borucki 1996; Giribet et al. 1999; Edgecombe and Giribet 2004; Murienne et al. 2010; Fernández et al. 2014b). The three families of the order include a total of ca. 95 species. More striking is the case of Craterostigmomorpha, with only two extant species (see Results and Discussion). These two orders originated more than 400 Mya, thus exhibiting very long branches prior to their respective diversification. In millipedes, the order Polyxenida includes only 89 species, and it is universally recognized as sister group to the other Diplopoda. As the basalmost groups of both centipedes and millipedes involve long branches, we also explored the effect of LBA in a fourth lineage with long branches and poor taxon representation, Symphyla, which were recovered as sister group to Polyxenida in some of our analyses (see Results and Discussion). To determine whether or not long branch attraction was affecting tree topology, we applied the SAW method of Siddall and Whiting (1999). This method is based on the removal of taxa in a pair suspected to be affected by LBA, one at a time. If after re-running the analysis either of the taxa appears at different branch points in the absence of the other, LBA is postulated. To understand the potential effect of LBA in our data set between (a) Symphyla and Polyxenida, and (b) Scutigeromorpha and Craterostigmomorpha, we pruned one taxon at a time in each case from the most complete matrix, supermatrix III (123 genes), and re-ran ExaBayes and PhyML_PCMA, as described above.

As rogue taxa can frequently have a negative impact on topology or in a bootstrap analysis (see a review in Goloboff and Szumik 2015), we also explored the presence of putative wild card taxa in our data set with RogueNaRok (Aberer et al. 2013), with the aim of identifying a set of taxa that, if pruned from the underlying bootstrap trees, yielded a reduced consensus tree containing additional bipartitions or increased support values.

### Morphological data

Morphological characters for the set of species for which transcriptomes were available or newly generated were coded, principally drawing on existing datasets. These include characters bearing on myriapod phylogeny (Rota-Stabelli et al. 2011), higher level chilopod phylogeny (Edgecombe and Giribet 2004; Murienne et al. 2010), diplopod phylogeny (Blanke and Wesener 2014), scutigeromorphs (Edgecombe and Giribet 2006; Koch and Edgecombe 2006), scolopendromorphs (Vahtera et al. 2013), and geophilomorphs (Koch and Edgecombe 2012; Bonato et al. 2014a). Sources for morphological information used for coding fossil terminals is detailed below (“Sources of coding and dates for fossils”). The dataset is available in nexus format as Morphobank Project P2216, “The myriapod tree of life” (www.morphobank.org/index.php/MyProjects/List/select/project_id/2216).

The morphology dataset consists of 232 characters, of which characters 57, 72, 86, 101 and 109 were scored as ordered/additive. These data were analyzed using parsimony as an optimality criterion in TNT (Goloboff et al. 2008) with equal character weights. A heuristic search used 10,000 random stepwise addition sequences saving 100 trees per replicate, with TBR branch swapping. Since our goal was not to produce a phylogenetic hypothesis derived exclusively from morphological data, no further analyses, such as those using implied weighting, were conducted.

## Divergence time inference

In order to infer divergence time in the myriapod phylogeny, we included seven Palaeozoic and Mesozoic fossils in our data set: four centipedes (*Crussolum* sp., *Devonobius delta, Mazoscolopendra richardsoni* and *Kachinophilus pereirai*) and three millipedes (*Cowiedesmus eroticopodus, Archidesmus macnicoli* and *Gaspestria genselorum*). In addition, we included two outgroup fossils, a crustacean (*Rehbachiella kinnekullensis*) and a scorpion (*Proscorpius osborni*). Absolute dates follow the International Stratigraphic Chart v 2014/02.

### a) Sources of codings and dates for fossils

*<u>Crussolum</u>* <u>Shear in Shear et al., 1998</u>: The scutigeromorph genus *Crussolum* includes one formally named species, *C. crusseratum* Shear in Shear et al. (1998), known from isolated and fragmentary legs from the Middle Devonian Gilboa locality, Schoharie County, New York State. Fossils come from the upper part of the Panther Mountain Formation, dated to the Tioughniogan regional Stage, Givetian in the global time scale. Palynomorphs are consistent with a Givetian age (Richardson et al. 1993). Legs and the forcipular segment of *Crussolum* sp. have been documented from the Windyfield Chert of the Dryden Flags Formation, Aberdeenshire, Scotland, by Anderson and Trewin (2003). An antenna was ascribed to this species by Anderson and Trewin (2003) but has not been used for coding because a hexapod identity cannot be ruled out. Spore assemblages of the Windyfield and stratigraphically underlying Rhynie Chert are dated to the early but not earliest Pragian to early (earliest?) Emsian (*polygonalis-emsiensis* Spore Assemblage Biozone) (Parry et al. 2011). Radiometric dating of the underlying Milton of Noth Andesite at ca 411 Ma (Parry et al. 2011, 2013) has been subject to a dispute over its temporal relationship to hot spring activity associated with the cherts (Mark et al. 2011; Mark et al. 2013) and predates the biostratigraphic dating of the Rhynie Chert relative to the global dating of the base of the Pragian Stage. We apply a date of the 407.6 Ma to the Rhynie and Windyfield cherts, using the Pragian-Emsian boundary as a reference. The coding for *Crussolum* draws mostly on the Windyfield Chert material, and accordingly is dated at 407.6 Ma. The oldest samples of *Crussolum* consist of isolated legs from Ludford Lane in England (Shear et al., 1998), sourced from a horizon 0.15-0.20 m above the base of the Ludlow Bone Bed Member of the Downtown Castle Sandstone Formation. The Ludlow Bone Bed Member is earliest Přidolí in age (Jeram et al., 1990).

*<u>Devonobius delta</u>* <u>Shear and Bonamo, 1988</u>: Described as the type species of a monotypic centipede order Devonobiomorpha, *Devonobius delta* is coded based on figured material of Shear and Bonamo (1988), all from Gilboa, New York. Stratigraphic details are identical to *Crussolum crusseratum* (see above). A minimum date for the end of the Givetian/base of the Frasnian is applied (382.7 Ma).

*<u>Mazoscolopendra richardsoni</u>* <u>Mundel, 1979</u>: Codings for this scolopendromorph centipede are based on descriptions and figures by Mundel (1979) and Haug et al. (2014), and personal observation by GDE of material in the Field Museum. Specimens are derived from the Francis Creek Shale Member of the Carbondale Formation, Mazon Creek, Illinois, of Westphalian D age (Shabica and Hay 1997). In the global time scale, this falls within the Moscovian Stage. A date for the top of the Moscovian/base of the Kasimovian is applied (307.0 Ma).

*<u>Kachinophilus pereirai</u>* <u>Bonato et al., 2014</u>: Coding of this geophilomorph centipede is based on the original description and direct study of the type material (Bonato et al. 2014b). U-Pd dating of zircons in rock matrix in “burmite” (Burmese amber) establishes a maximum age of 98.79 +/− 0.62Ma (early Cenomanian) for amber inclusions (Shi et al. 2012).

*<u>Cowiedesmus eroticopodus</u>* <u>Wilson and Anderson, 2004:</u> This species is one of three co-occurring species assigned to Archipolypoda, an extinct superordinal level grouping identified as Chilognatha (Wilson and Anderson, 2004). These represent the oldest body fossils of Diplopoda. Coding is based on the original description and figures of the holotype by Wilson and Anderson (2004) and personal examination by GDE of the specimen in the Australian Museum. This species occurs in the *Dictyocaris* Member of the Cowie Formation in the Stonehaven Group, Stonehaven, Scotland. As summarized by Wilson and Anderson (2004), associated palynomorphs date the unit to the late Wenlock to early Ludlow (Silurian). A minimum date for the early Ludlow uses the base of the Ludfordian Stage (425.6 Ma).

*<u>Archidesmus macnicoli</u>* <u>Peach, 1882:</u> Selected to supplement *Cowiedesmus* in providing morphological characters for Archipolypoda, coding is based on the revision of this species by Wilson and Anderson (2004). Specimens figured therein come from the Dundee Formation in the Arbuthnott Group, Tillywhandlung Quarry, near Forfar, Scotland. Miospores date the unit to the *micrornatus-newportensis* biozone, of Lochkovian (Lower Devonian) age (Wilson and Anderson 2004). A minimum date is applied to Lochkovian-Pragian boundary (410.8 Ma).

*<u>Gaspestria genselorum</u>* <u>Wilson, 2006:</u> Coding of this diplopod is based on the original description and published illustrations by Wilson (2006), who interpreted *G. genselorum* as the oldest juliformian, and new photographs of the types provided by R. Miller. Wilson (2006) reviewed the plant megafossil and spore constraints on the age of the two units in which the species occurs, the Cap-aux-Ocs Member of the Battery Point Formation (Gaspé, Québec) and the Campbelltown Formation (New Brunswick). Spores at the Gaspé site are assigned to the *sextantii* Subzone of the *annulatus-lindlarensis* Zone and those of the New Brunswick site to the *Grundiospora* Subzone of the *annulatus-lindlarensis* Zone. Both are of late Emsian age. A minimum date of 393.3 Ma is accordingly applied, constrained by the Emsian-Eifelian boundary.

*<u>Proscorpius osborni</u>* <u>(Whitfield, 1885)</u>: This species was selected as being an anatomically well-understood Silurian scorpion (Whitfield 1885), using the revision by Dunlop et al. (2008) for coding. Formerly known as the ‘Bertie Waterlime’, the stratigraphic unit from which the material was sourced is the Phelps Member of the Fiddlers Green Formation, in the Bertie Group, New York State. This unit is dated to the Přídolí Series of the Silurian (Dunlop et al. 2008), and assigned a minimal age of 419.2 Ma (dating the Přídolí-Lochkovian boundary).

*<u>Rehbachiella kinnekullensis</u>* <u>Müller, 1983</u>: The monographic revision by Walossek (1993) was used for coding. Material documented therein was derived from four localities of the late Cambrian *Orsten* in southern Sweden. These are assigned to the *Agnostus pisiformis* Zone and, less commonly, to the overlying *Olenus gibbosus* Zone (late Guzhangian and earliest Paibian, respectively: Nielsen et al. (2014)). Because some of the material is demonstrably Guzhangian, a minimal date is applied to the bases of the Furongian and Paibian, 497 Ma.

### b) Divergence time inference through node calibration

Temporal data for the fossils listed above provide constraints on divergence dates. The resolution of *Crussolum* as stem-group Scutigeromorpha in our morphological cladogram (Fig. 3; Fig. S3) constrains the split of Scutigeromorpha and Pleurostigmophora, i.e., crown-group Chilopoda, to at least 419.2 Ma, drawing on the Ludlow Bone Bed material of *Crussolum. Devonobius delta* is resolved in our morphological analysis as sister group of Epimorpha but an alternative affinity to *Craterostigmus* has been suggested (Borucki 1996). To allow for uncertainty over the phylogenetic position of *Devonobius* and *Craterostigmus,* the fossil was conservatively used only to constrain crown-group Pleurostigmophora (minimum of 382.7 Ma based on *Devonobius*). Crown-group Epimorpha is constrained to a minimum of 306 Ma by the oldest scolopendromorph *Mazoscolopendra richardsoni.* That species is resolved as total-group Scolopendromorpha based on our morphological analysis (Fig. S3). *Kachinophilus pereirai,* originally assigned to Geophilidae (Bonato et al. 2014b), is resolved based on morphology in a clade with extant geophilids (Fig. S3). As such, it constrains crown-group Adesmata, i.e., the split between Mecistocephalidae and remaining Geophilomorpha, to at least 98.79 Ma. Crown-group Chilognatha is constrained by the occurrence of *Cowiedesmus eroticopodus,* resolved in our morphological analysis as total-group Helminthomorpha (425.6 Ma). *Rehbachiella kinnekullensis* has been interpreted as stem-group Anostraca, i.e., crown-group Branchiopoda (Walozsek, 1993, reference phylogeny Fig. 41 therein), or as stem-group Branchiopoda (Olesen 2009; reference phylogeny in Fig. 14 therein). Our morphological data agree with branchiopod affinities in resolving it as more closely related to *Daphnia* than to *Calanus.* This provides a minimum date for crown-group Altocrustacea (sensu Regier et al. 2010) to 497 Ma. Because of uncertainty with regards to the interrelationships between major pancrustacean groups (including Branchiopoda, Copepoda and Insecta, from which the three exemplars used here were sampled) this date is conservatively applied only to crown-group Pancrustacea. *Proscorpius osborni* is resolved by our morphological data dataset as closest relative of the extant scorpion exemplar *Centruroides,* setting a constraint on the split of Scorpiones from Tetrapulmonata. The relevant crown-group based on our taxon sampling is Arachnopulmonata, dated to at least 419.2 Ma.

**Figure 3.**
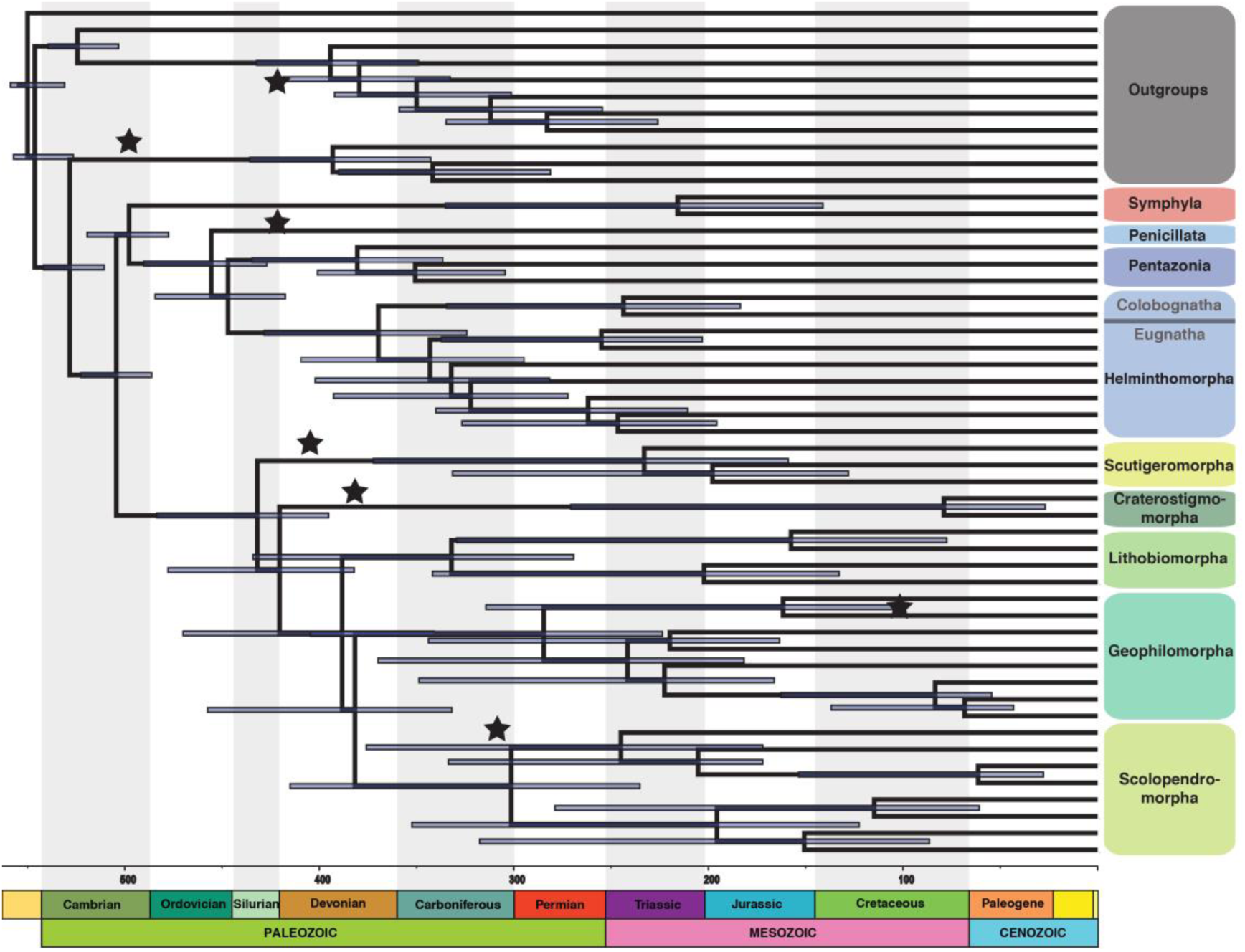
Chronogram of myriapod evolution for the 123-gene data set with 95% highest posterior density (HPD) bar for the dating under the uncorrelated gamma model. Nodes that were calibrated with fossils are indicated with a star placed at the age of the fossil.

The split between Onychophora and Arthropoda was dated between 528 million years (the minimum age for Arthropoda used by Lee et al. 2013 on the basis of the earliest *Rusophycus* traces in the early Fortunian, dating the base of Cambrian Stage 2) and 558 million years. The latter was used as the root of Panarthropoda (Lee et al. 2013) based on dating of White Sea Vendian strata that contain the oldest plausible bilaterian fossils.

Divergence dates were estimated using the Bayesian relaxed molecular clock approach as implemented in PhyloBayes v.3.3f (Lartillot et al. 2013). An auto-correlated model of clock relaxation was applied as it has been shown to provide a significantly better fit than uncorrelated models on phylogenomic datasets (Lepage et al. 2007; Rehm et al. 2011). The calibration constraints specified above were used with soft bounds (Yang and Rannala 2006) under a birth-death prior in PhyloBayes, this strategy having been found to provide the best compromise for dating estimates (Inoue et al. 2010). Two independent MCMC chains were run for 5,000-7,000 cycles, sampling posterior rates and dates every 10 cycles. The initial 25% were discarded as burn-in. Posterior estimates of divergence dates were then computed from the remaining samples of each chain.

## RESULTS AND DISCUSSION

### Phylogenetic relationships of Myriapoda

All analyses conducted, regardless of methodology or data set, recover monophyly of Myriapoda, Diplopoda, Chilopoda and Symphyla, but the interrelationships among the myriapod classes are less stable (Fig. 2). With regards to the position of Symphyla, two alternatives are found: Symphyla either unite with Diplopoda as predicted by the Progoneata hypothesis or Chilopoda and Diplopoda unite as a clade to the exclusion of Symphyla. The latter result was also found in many of the analyses by Rehm et al. (2014). A chilopod-diplopod clade has not been anticipated morphologically, though certain morphological characters fit such a grouping. For example, chilopods and diplopods share a series of imbricated comb lamellae on the mandibles that are lacking in symphylans and pauropods (Edgecombe and Giribet 2002). This character was proposed as a potential myriapod autapomorphy but the standard phylogenetic hypotheses forced reversals/losses in pauropods and symphylans. Under one of our two scenarios, although in the absence of pauropods, this character would be interpreted as a synapomorphy of Chilopoda + Diplopoda. In the analyses designed to account for LBA, such as the SAW analysis (Fig. S4) with Polyxenida removed and the PhyloBayes analysis with the CAT model of heterogeneity, Symphyla was recovered as sister group to the clade Diplopoda + Chilopoda, suggesting that Symphyla could have been attracted to Diplopoda due to the long branch of Polyxenida. It need be noted, however, that a position of Symphyla closer to Diplopoda than to Chilopoda has been endorsed morphologically (Progoneata hypothesis) (Dohle 1980; Edgecombe 2004, 2011).

The 4-gene analysis including the sequences available for a pauropod recovered Pauropoda as sister group to Symphyla with posterior probability = 0.99. However, the relationships with the other myriapod groups remained unsupported, and the pauropod-symphylan group (=Edafopoda) is observed to be associated with anomalous relationships between orders of both Chilopoda and Diplopoda (Fig. S1). A denser taxon sampling of the two lesser myriapod groups, Symphyla (e.g., including representatives of the other family, Scolopendrellidae) and Pauropoda, would be definitely key to resolve the backbone of the myriapod tree with greater confidence.

*Diplopoda:* Within Diplopoda, all analyses (apart from the one including a pauropod; Fig. S4) support a sister group relationship between Penicillata (sampled by *Eudigraphis*) and all other millipedes, the latter forming the traditional clade Chilognatha. Chilognatha is united by a calcified cuticle and the male actively transferring sperm to the female genital opening, among other apomorphic characters (Enghoff 1984; Blanke and Wesener 2014). The fundamental division within Chilognatha likewise corresponds to traditional morphology-based systematics, including a sister group relationship between Pentazonia (generally pill millipedes) and Helminthomorpha (long-bodied millipedes).

Relationships within Pentazonia depart from the standard morphological hypothesis. The latter usually unites Glomerida and Sphaerotheriida as sister groups (collectively forming a clade of pill millipedes named Oniscomorpha) to the exclusion of Glomeridesmida. In contrast to this, however, our analyses resolve Sphaerotheriida (*Cyliosoma*) as sister group to Glomerida (*Glomeris*) + Glomeridesmida (*Glomeridesmus*). The only previous molecular studies to include all three orders of Pentazonia are the three-gene analysis of Regier et al. (2005) and the same dataset combined with morphological data by Sierwald and Bond (2007). As in our analyses, these yielded an “unexpected” (Regier et al. 2005: p. 155) grouping of Glomeridesmida and Glomerida to the exclusion of Sphaerotheriida. Monophyly of Oniscomorpha has been defended morphologically based on the presence of a collum much smaller than the following tergites, the second tergite being much larger than following tergites, and a the lack of coxal pouches on legs (Blanke and Wesener 2014). The status of these characters as synapomorphies is disputed by our trees, and instead are interpreted as plesiomorphies of Pentazonia.

In Helminthomorpha, the traditional clades Colobognatha and Eugnatha are each monophyletic, as found in prior analyses of largely the same set of millipede species (Brewer and Bond, 2013), as well as by sampling different exemplars for three nuclear coding genes (Miyazawa et al. 2014). The colobognathan terminals are *Brachycybe* (order Platydesmida) and *Petaserpes* (order Polyzoniida); transcriptomic data remain unavailable for two other colobognathan orders, Siphonocryptida and Siphonophorida. Colobognatha and Eugnatha were originally proposed in the classical era of myriapod systematics (Attems 1926; Verhoeff 1928), but were discarded in the influential millipede classification by Hoffman (1980), who considered that the characteristic reduced mouthparts of Colobognatha were prone to convergence. They have, however, consistently been endorsed in cladistic analyses (Enghoff 1984; Sierwald et al. 2003; Blanke and Wesener 2014). As likewise found by Brewer and Bond (2013), Eugnatha includes two groups that have a long history of usage in morphology-based systematics. One of these unites Chordeumatida (sampled by *Cleidogona*) and Callipodida (sampled by *Abacion*), which are classified together as Coelochaeta. Another clade consists of the ring-forming millipedes, Juliformia, which unites the sampled members of Julida (*Cylindroiulus*), Spirostreptida (*Cambala*) and Spirobolida (*Narceus*). Our analyses variably resolve the three-taxon problem of the juliform orders with either Julida or Spirobolida as sister group of Spirostreptida. Both of these conflict with the weak morphological support for Spirobolida and Julida as closest relatives (Blanke and Wesener 2014), but have been found in other molecular studies. A Julida + Spirostreptida clade was supported by three nuclear protein-coding genes (Miyazawa et al. 2014), whereas nuclear ribosomal genes (Cong et al. 2009) and mitochondrial gene rearrangements and amino acid sequences (Woo et al. 2007) supported a Spirobolida + Spirostreptida clade. The relationships of the juliform orders need be regarded as an open question for the time being.

Two of the most contentious issues in millipede systematics involve the relationships of Polydesmida and of Stemmiulida. From the perspective of morphology, Polydesmida has been considered the sister group of Juliformia as a “ring-forming” clade (Enghoff et al. 1993) or as sister group to a putative clade named Nematophora (Sierwald et al. 2003; Blanke and Wesener 2014). The latter group, named for the shared presence of preanal spinnerets, traditionally includes Stemmiulida, Callipodida and Chordeumatida (Enghoff 1984; Sierwald et al. 2003). Spinnerets are, however, more widely shared by Polydesmida (Shear 2008) and this character has been cited as synapomorphic for a clade containing Polydesmida + Nematophora (Blanke and Wesener 2014). Our results, like those of Brewer and Bond (2013), do not retrieve monophyly of Nematophora in its traditional guise: Chordeumatida + Callipodida do not unite with Stemmiulida, the latter being resolved either with Polydesmida or with Juliformia. Optimization on the topology of Brewer and Bond (2013) as well as those obtained here suggests that spinnerets are an autapomorphy of Eugnatha as a whole and were lost in Juliformia.

*Chilopoda:* Half of the analyses retrieved the widely accepted division of Chilopoda into Notostigmophora (Scutigeromorpha) and Pleurostigmophora (the other four orders). However, in the other half Scutigeromorpha and Craterostigmomorpha were recovered as a clade. This result has never been advocated morphologically. Both clades are invariantly at the base of the tree, as recovered in all the SAW analyses (Fig. S6). However, their interrelationships remain unresolved. A possible cause of this phylogenetic conundrum could be the low diversity of Craterostigmomorpha, which comprises only two extant species. Within Pleurostigmophora, the Amalpighiata hypothesis (Craterostigmus sister group of all other pleurostigmophorans; Fernández et al. 2014b) withstood all procedures designed to target systematic error, such as increased gene sampling and the exploration of complex evolutionary models, and was favored over the Phylactometria hypothesis (Lithobiomorpha sister group of all other pleurostigmophorans; Edgecombe and Giribet 2004). The relationships between Lithobiomorpha and Epimorpha (= Scolopendromorpha and Geophilomorpha) remained elusive, as several analyses recovered Scolopendromorpha as sister group to the rest of this clade, defying nearly all previous morphological and molecular evidence. Notably, all the matrices recovering this topology include the highest gene occupancy (supermatrices III-V) at the expense of numbers of genes (see below for further details).

All chilopod orders are well supported and relationships within each of the large orders correspond to previous phylogenetic hypotheses, and for most to traditional taxonomy. The three families of Scutigeromorpha resolve with Pselliodidae (*Sphendononema*) as sister group to Scutigerinidae (*Scutigerina*) and Scutigeridae (*Scutigera*). The same topology has consistently been inferred using targeted sequencing of a few loci (Edgecombe and Giribet 2006, 2009; Butler et al. 2010; Giribet and Edgecombe 2013), but contrasts with the morphological placement of Scutigerinidae as sister group to Pselliodidae and Scutigeridae (Koch and Edgecombe 2006). Lithobiomorpha comprises two clades that correspond to its two families, Henicopidae (*Anopsobius* and *Paralamyctes*) and Lithobiidae (*Lithobius* and *Eupolybothrus*). Scolopendromorpha consists of a blind clade and an ocellate clade. The blind clade, supported in all analyses, unites members of Cryptopidae, Scolopocryptopidae and Plutoniumidae whereas the ocellate clade is the family Scolopendridae. The number of events of eye loss in Scolopendromorpha had been a subject of some debate (Vahtera et al. 2012) but the transcriptomic data strongly corroborate weaker signal from targeted gene sequencing (Vahtera et al. 2013) and morphology (Koch et al. 2009; Koch et al. 2010) for blindness in the common ancestor of Cryptopidae and Scolopocryptopidae + Plutoniumidae. All analyses herein resolve Scolopocryptopidae as paraphyletic, *Scolopocryptops* being more closely related to *Theatops* (Plutoniumidae) than to *Newportia.* This result implies that the shared presence of 23 leg-bearing trunk segments in Scolopocryptopinae (*Scolopocryptops*) and Newportiinae (*Newportia*) is either acquired convergently from a 21-segmented ancestor or is homologous but was reversed (to 21 segments) in Plutoniumidae. Monophyletic Scolopendridae consists of Otostigminae (*Alipes* and *Rhysida*) and Scolopendrinae (*Akymnopellis* and *Scolopendropsis*), precisely mirroring classical and current taxonomy. Geophilomorpha also resolves along traditional lines into Placodesmata (Mecistocephalidae: *Mecistocephalus* and *Tygarrup*) and Adesmata, a clade composed of all other geophilomorphs. Resolution within Adesmata conflicts with the most recent analysis of geophilomorph relationships (Bonato et al. 2014); they found a clade containing Oryidae, Himantariidae and Schendylidae (Himantarioidea), while our analyses likewise recover a Himantariidae + Schendylidae clade but place *Notiphilides* (Oryidae) as sister group to Geophilidae, a result obtained for every analysis of each data set. However, we were not able to fully test the monophyly of Geophiloidea, as Zelanophilidae and Gonibregmatidae are not represented in our analyses. Our three Geophilidae (representatives of Geophilidae and two recently synonymized families Dignathodontidae and Linotaeniidae) constitute a well-supported and stable clade.

#### Tempo of diversification in myriapods

The dated phylogeny for the 123 gene matrix reflects a diversification of Myriapoda in the early and middle Cambrian, supporting a Cambrian diversification event, as suggested recently for this lineage (Rota-Stabelli et al. 2013). This dating continues to substantially predate body fossil or even trace fossil evidence for Myriapoda, the oldest trackways that can be ascribed to myriapods with reasonable confidence being Late Ordovician in age (Johnson et al. 1994; Wilson 2006). Diversification of Diplopoda is inferred to occur in the late Cambrian to mid Silurian, and the diversification of Chilopoda in the Early Ordovician to Middle Devonian (Fig. 3). The diversification times of the millipede and centipede orders included in our analyses are thus generally congruent with the timing inferred in previous studies (Meusemann et al. 2010; Rehm et al. 2011; Brewer and Bond 2013; Fernández et al. 2014b), and do not greatly predate first occurrences of major chilopod and diplopod lineages in the fossil record. On the other hand, an estimated Cambrian diversification of myriapods implies a large gap in the fossil record. Fossil candidates for stem-group Myriapoda in marine, freshwater or terrestrial sediments of Cambrian age remain unknown or unidentified (Edgecombe 2004; Shear and Edgecombe 2010). Nonetheless, the occurrence of crown-group Pancrustacea as early as Cambrian Stage 3 (Edgecombe and Legg 2014) predicts a ghost lineage for Myriapoda to at least that time, ca 517 Ma.

In centipedes, the additional transcriptomes with respect to previous studies (Fernández et al. 2014b) resulted in slightly older dates, but it did not affect their origin. For instance, the latter study inferred the diversification of Pleurostigmophora in the Middle Ordovician to Early Carboniferous, the diversification of Amalpighiata in the Silurian-Carboniferous, and the diversification of Epimorpha from the Early Devonian to the early Permian. In this study, the recovered dates ranged from the Early Ordovician to the Middle Devonian in Pleurostigmophora, from the Middle Ordovician to the Early Carboniferous in Amalpighiata, and Epimorpha from the Late Devonian to the mid Carboniferous.

The controversy over the systematic position of *Craterostigmus* (i.e., whether Phylactometria or Amalpighiata is monophyletic) relates to lineages that have ancient, rapidly diverging stem groups but relatively shallow crown groups, conditions expected to contribute to a difficult phylogenetic problem. Despite a divergence from other living chilopod orders by the Late Silurian, the two extant species of *Craterostigmus* have a mean date for their divergence from each other in the Cretaceous, and these species are almost indistinguishable morphologically (Edgecombe and Giribet 2008). Likewise, Scutigeromorpha, with which *Craterostigmus* groups in various analyses, has a stem dating to the Silurian but the deepest split between its three extant families has a mean date in the Triassic. Although the group is also conservative morphologically, we have maximized phylogenetic diversity here for a clade that has been well resolved phylogenetically and which increased its diversification rate around 100 million years ago (Giribet and Edgecombe 2013). This is effectively like Craterostigmomorpha, where all the extant diversity is comparatively recent and old branches are unrepresented in our analyses because of extinction.

#### Using morphology to test phylogenomic hypothesis

Results herein demonstrate that most systematic hypotheses for Myriapoda are strongly supported across the explored sets of genes, optimality criteria and programs. For example, the deep interrelationships within the four large centipede orders are stable across all analyses (Fig. 2). These relationships also show a high degree of congruence with morphological trees and morphology-based classifications. In the case of, for example, Scolopendromorpha, we can conclude that a high degree of confidence can be placed in the results.

Where hypotheses vary according to different methods or data partitions, in some cases morphology provides a strong arbiter for evaluating the rival hypotheses. For example, the 123-gene analyses typically recover Lithobiomorpha as sister group of Geophilomorpha, whereas the larger gene samples generally group Geophilomorpha with Scolopendromorpha. The former hypothesis has, to our knowledge, only been recovered in a single, non-numerical phylogenetic analysis, based on a single character of sperm structure (Jamieson 1986). In contrast, a geophilomorph-scolopendromorph clade – the classical Epimorpha – receives morphological support from the perspectives of development, behavior, external morphology and internal anatomy (eight autapomorphies listed by Edgecombe 2011). This ability of morphology to select between rival hypotheses that are each based on vast pools of data affirms the continued relevance of morphological characters in phylogenetics (e.g., Giribet 2015; Lee and Palci 2015; Wanninger 2015) and permit delving into the possible reasons for the discordance between the different analyses.

Furthermore, access to a morphological dataset allowed us to establish the position of fossil terminals for node calibration using the exact same set of extant terminals used in phylogenomic analyses (as in Sharma and Giribet 2014). Most recent justification for the necessity of morphological data in molecular dating has focused on its indispensability for so-called tip dating or total evidence dating (e.g., Giribet 2015; Lee and Palci 2015; Pyron 2015). However, even in the standard node calibration approach as employed here, the position of fossils can be determined in the context of the precise taxonomic sample used in other parts of the study, rather than cobbling together justifications for node calibrations from external (or possibly mixed) sources.

#### The effect of confounding factors in phylogenomic reconstruction: matrices with high taxon occupancy recovered the ‘wrong’ topologies

Assessing the effect of missing data on phylogenetic inference has received substantial attention from phylogeneticists over many years, but unambiguous conclusions have been evasive. Properties of the data sets themselves, such as different rates of evolution and compositional heterogeneity, can have a strong influence on the accuracy of phylogenetic inferences, irrespective of the amount or the pattern of missing data. The combined effect of these factors complicates the problem, and no study has dissected out in depth the effect of each parameter independently. The different matrices constructed and the multitude of analyses conducted to address the potential effects of a diversity of confounding factors in phylogenomic inference (missing data, compositional heterogeneity, heterotachy, differential evolutionary rate, levels of phylogenetic informativeness, model misspecification, concatenation and gene-tree incongruence) yield congruent results (see Fig. 2, Fig. S2, Fig. S3, Fig. S5, Fig. S6). The main three conflicting nodes (marked as 1, 2 and 3 in Fig. 2) are recovered in approximately half of the analyses, indicating that these artifactual/methodological explanations for the incongruence alone are unlikely to have driven the recovered topologies.

The potential impact of missing data on phylogenomic inference accuracy has been poorly investigated, particularly for higher-level (=deep and old) relationships, but gene completeness has often been advocated as the only justification for selecting genes. This has sometimes been used as a justification for using target-enrichment approaches to phylogeny versus ‘phylotranscriptomics’ (e.g., Lemmon and Lemmon 2013). In the past decades, several studies have investigated the effects of missing data in phylogenetic reconstruction (e.g., Maddison 1993; Norell et al. 1995; Norell and Wheeler 2003; Wiens 2003; Simmons 2012), with some recent studies focusing on phylogenomic data sets with hundreds or thousands of orthologous genes (e.g., Rokas et al. 2003; Philippe et al. 2004; Roure et al. 2013). These studies have suggested a negative (e.g., missing data have a deleterious effect on accuracy) or neutral (e.g., missing data *per se* do not directly affect inference) effect of the missing data. The negative effect is mainly due to a reduced detection of multiple substitutions, exacerbating systematic errors such as long-branch attraction (LBA), as incomplete species are less efficient in breaking long branches (Roure et al. 2013)--but that study demonstrated this using a matrix too small to resolve deep metazoan relationships. Perhaps for this reason other studies have shown that adding incomplete taxa is not deleterious *per se* as long as enough informative character states are present for each species, but analyzing too few complete characters could reduce accuracy because of a lack of phylogenetic signal (Wiens 2003; Philippe et al. 2004; Wiens 2006; Lemmon et al. 2009). In this study, we show for the first time that high matrix occupancy can lead to ‘incorrect’ inferences: all the different analyses conducted with the most complete matrix (supermatrix III) resulted in a topology of centipedes at odds with well-established phylogenetic hypotheses and with nearly all other analyses using more genes or selecting genes based on other criteria (informativeness, compositional homogeneity, etc.), and curiously it was the matrix that showed the highest level of conflict between individual gene trees (Fig. S6). These three contradicting ideas on the effect of missing data in phylogenetic inference hinder the experimental design in phylogenomics, and highlight the necessity of further exploring its effect in new data sets.

The conflicting nodes between centipede orders are unlikely to result from incomplete taxon sampling, as our data set includes all extant families of centipedes with the exception of two geophilomorph families (Zelanophilidae and Gonibregmatidae) and one monotypic family of Scolopendromorpha (Mimopidae). In this context, biological explanations for the incongruence, such as incomplete lineage sorting consistent with a scenario of rapid radiation, or lineage-specific high extinction, may need to be considered. In fact, Craterostigmomorpha, the taxon responsible for some of the major instability, comprises only two species in Tasmania and New Zealand, constituting an old lineage with depauperate extant diversity and conservative morphology, some of the conditions often used to refer to living fossils (Werth and Shear 2014).

### Concluding Remarks

In this study we explored the phylogeny of Myriapoda and the internal relationships of its largest clades, Diplopoda and Chilopoda, using published genomes and transcriptomes as well as novel transcriptomic data for 25 species. For this we constructed an array of data sets to independently optimize gene number, gene occupancy, phylogenetic informativeness, or gene conservation and analyzed them using different phylogenetic methods and evolutionary models. In addition we generated a morphological data matrix of 232 characters that was used to precisely place a set of fossils subsequently used for node calibration analyses. Our results largely corroborate those of previous analyses, especially with respect to the millipede ordinal relationships and added resolution to the centipede tree, sampled at the family-level. However, we identified a few conflicting nodes across analyses to discover that for the most part the matrices optimizing gene occupancy produced topologies at odds with morphology, development, prior molecular data sets, or most notably, our own analyses using matrices with larger numbers of genes (with lower average gene occupancy) or matrices optimizing phylogenetic informativeness and gene conservation. This calls for caution when selecting data sets based strictly on matrix completeness and adds further support to previous notions that a large diversity of genes, even to the detriment of matrix occupancy (contra Roure et al. 2013), may be a feasible solution to analyzing phylogenetic relationships among deep animal clades (e.g., Hejnol et al. 2009). Finally, our dating analysis continues to support results from prior studies in placing the diversification of Myriapoda in the Cambrian, prior to any recognizable myriapod fossil record, whereas the estimated diversification of both millipedes and centipedes is closer to the existing fossil record. Further refinement of the myriapod tree of life will require addition of pauropods, more symphylans, and a few missing lineages of centipedes and millipedes.

## Acknowledgements

We are grateful to many colleagues who provided tissues, among them Jesús Ballesteros-Chávez, Tony Barber, Beka Buckman, Amazonas Chagas, Jr., Savel Daniels, Ronald Jenner, Bob Kallal, Bob Mesibov, Ana Tourinho, and Varpu Vahtera. Fieldwork was supported by the Putnam Expedition Grants program of the MCZ and by NSF grant DEB #1144417 (Collaborative Research: ARTS: Taxonomy and systematics of selected Neotropical clades of arachnids) to G. Hormiga and G.G. Sequencing costs were mostly covered by internal MCZ and FAS funds. R.F. was supported by NSF grant DEB #1457539 (Collaborative Proposal: Phylogeny and diversification of the orb weaving spiders) to G. Hormiga and G.G.

---

**Figure S1.**
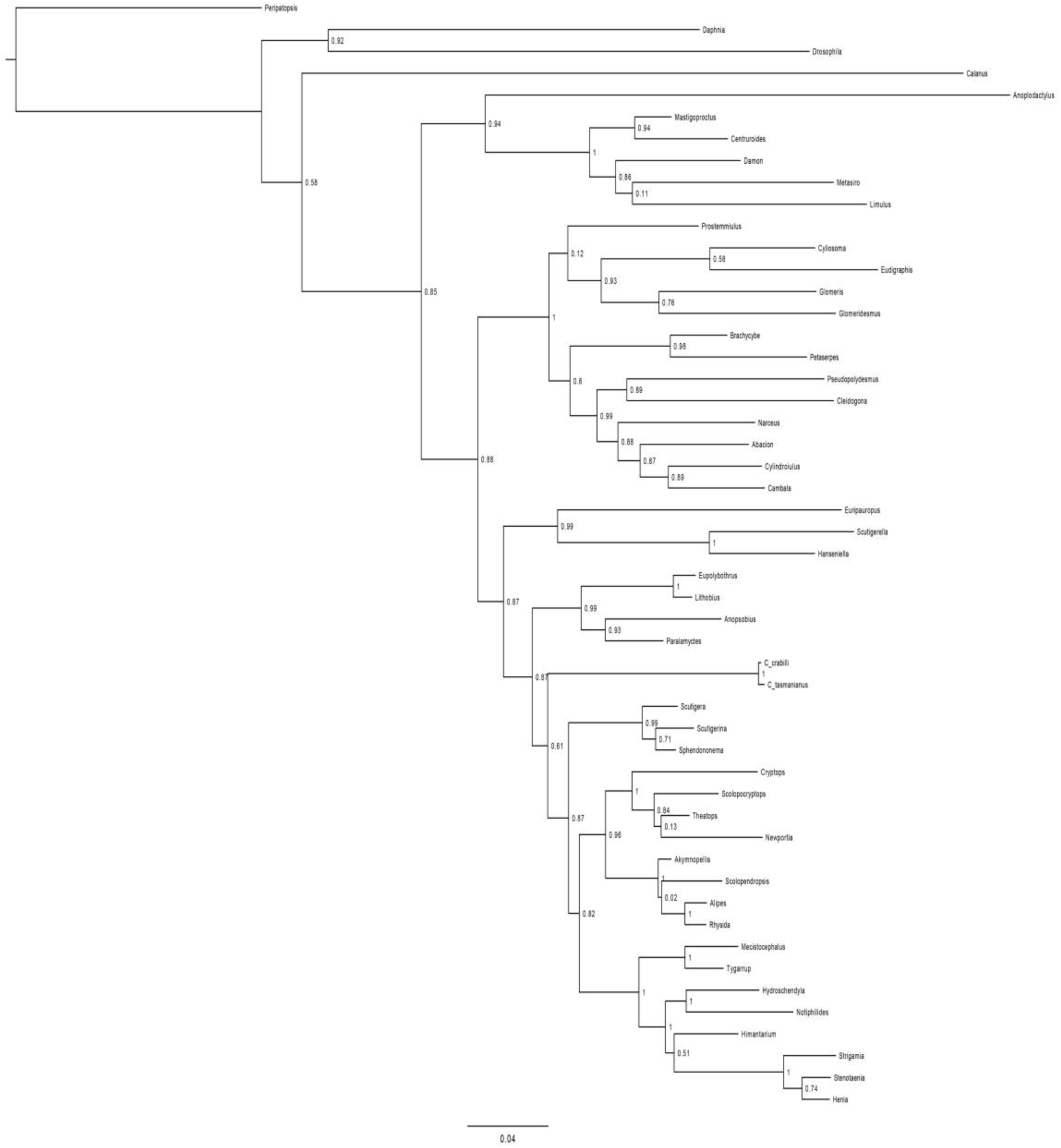
Phylogenetic hypothesis of the interrelationships of Pauropoda, Symphyla, Diplopoda and Chilopoda inferred from a 4-gene data set using ExaBayes.

**Figure S2.**
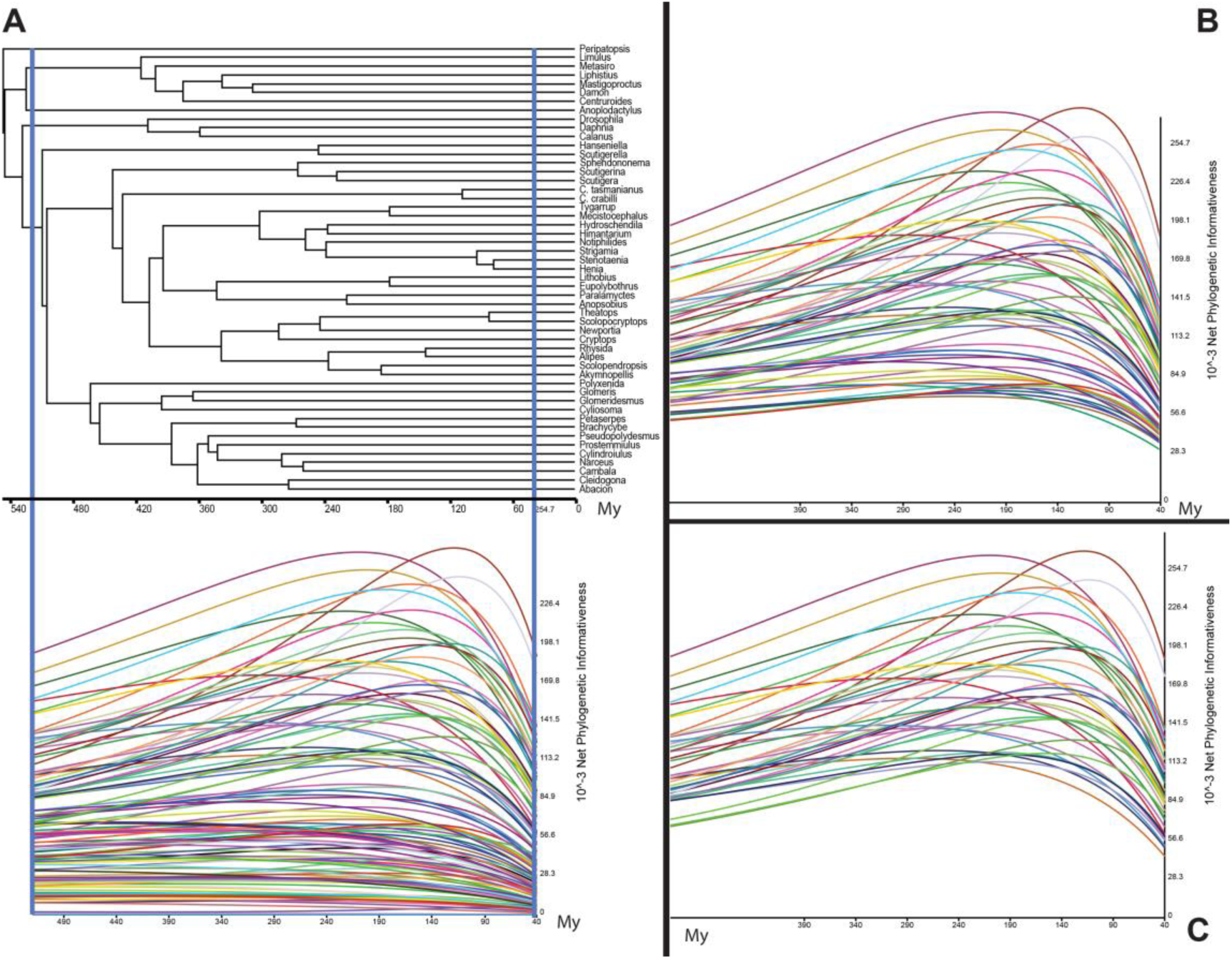
Phylogenetic informativeness profile of the genes included in supermatrices III (a), VI (b) and V (c) during the interval of time corresponding to the diversification of all myriapod clades (as delimited by blue lines in the ultrametric tree in (a)).

**Figure S3.**
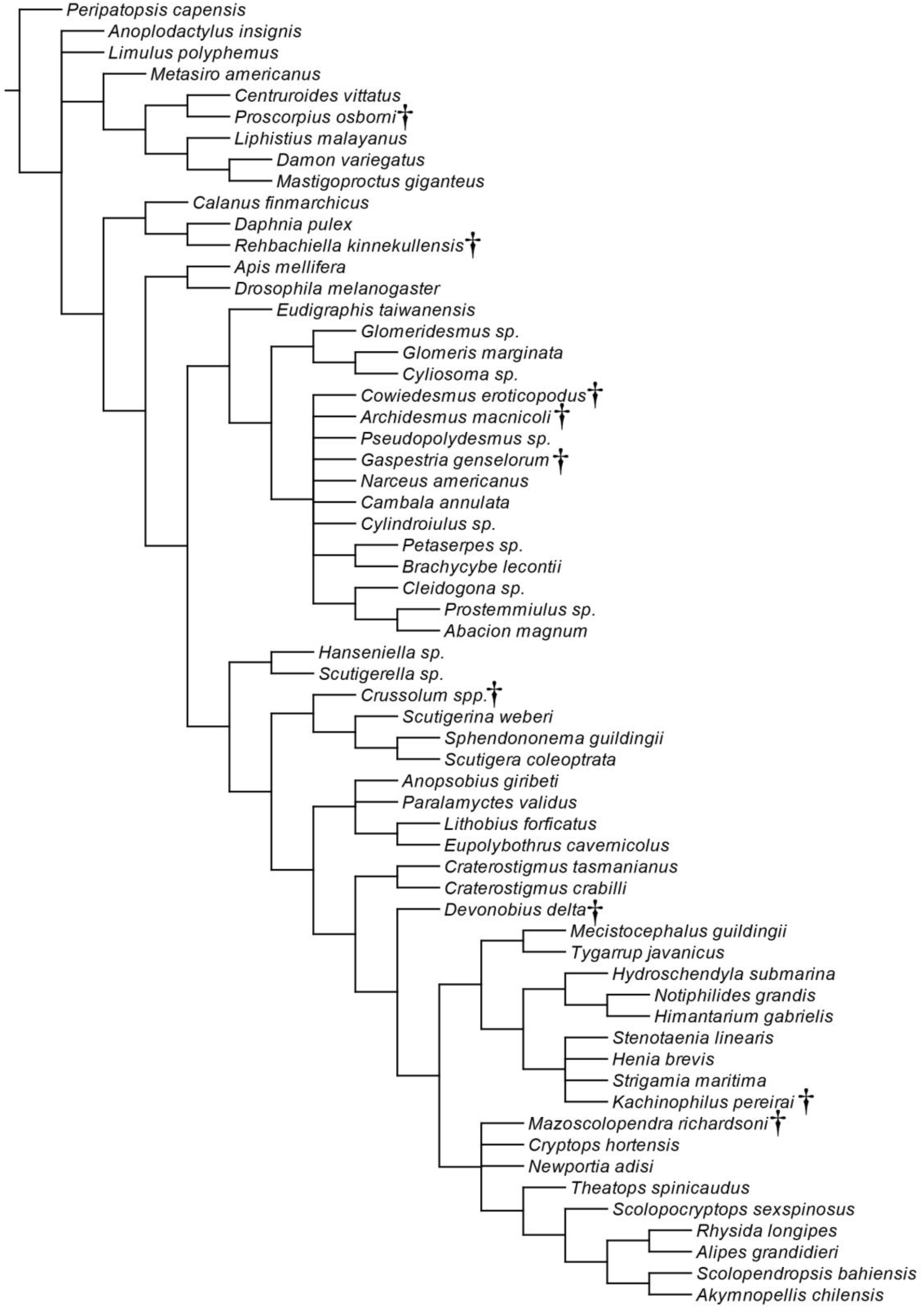
Phylogenetic hypothesis of Myriapoda based on 232 morphological characters coded for both extant and extinct species (see Material and Methods for further details). Strict consensus of 2515 shortest cladograms (420 steps). Fossil taxa are marked with a dagger symbol.

**Figure S4.**
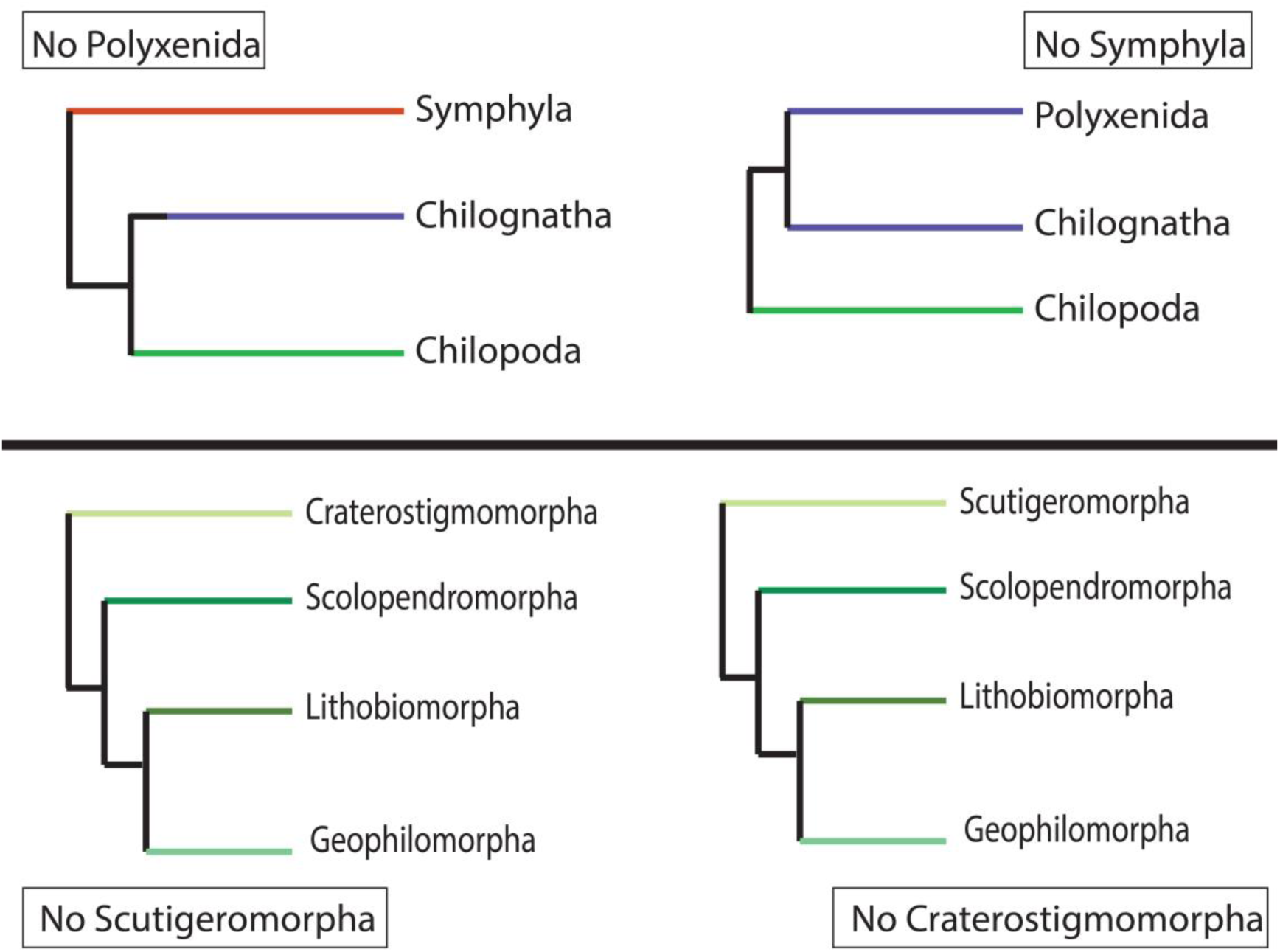
Topologies recovered in the SAW analyses to test for LBA between a) Symphyla and Polyxenida, and b) Scutigeromorpha and Craterostigmomorpha.

**Figure S5.**
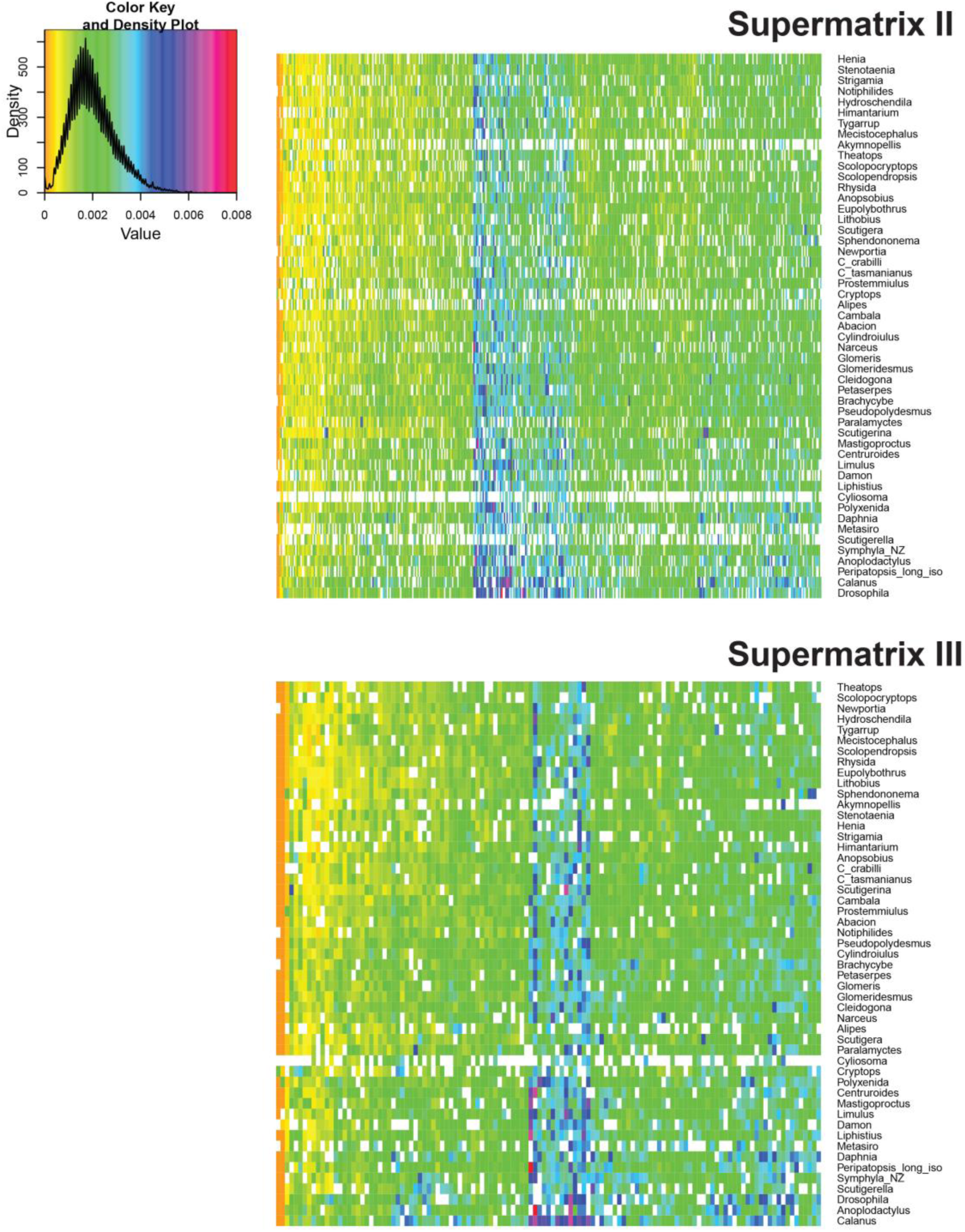
BaCoCa results showing compositional homogeneity values per taxon and per gene in supermatrices II and III (results in supermatrix I not shown due to a lack of resolution related to the high number of genes included in the matrix).

**Figure S6.**
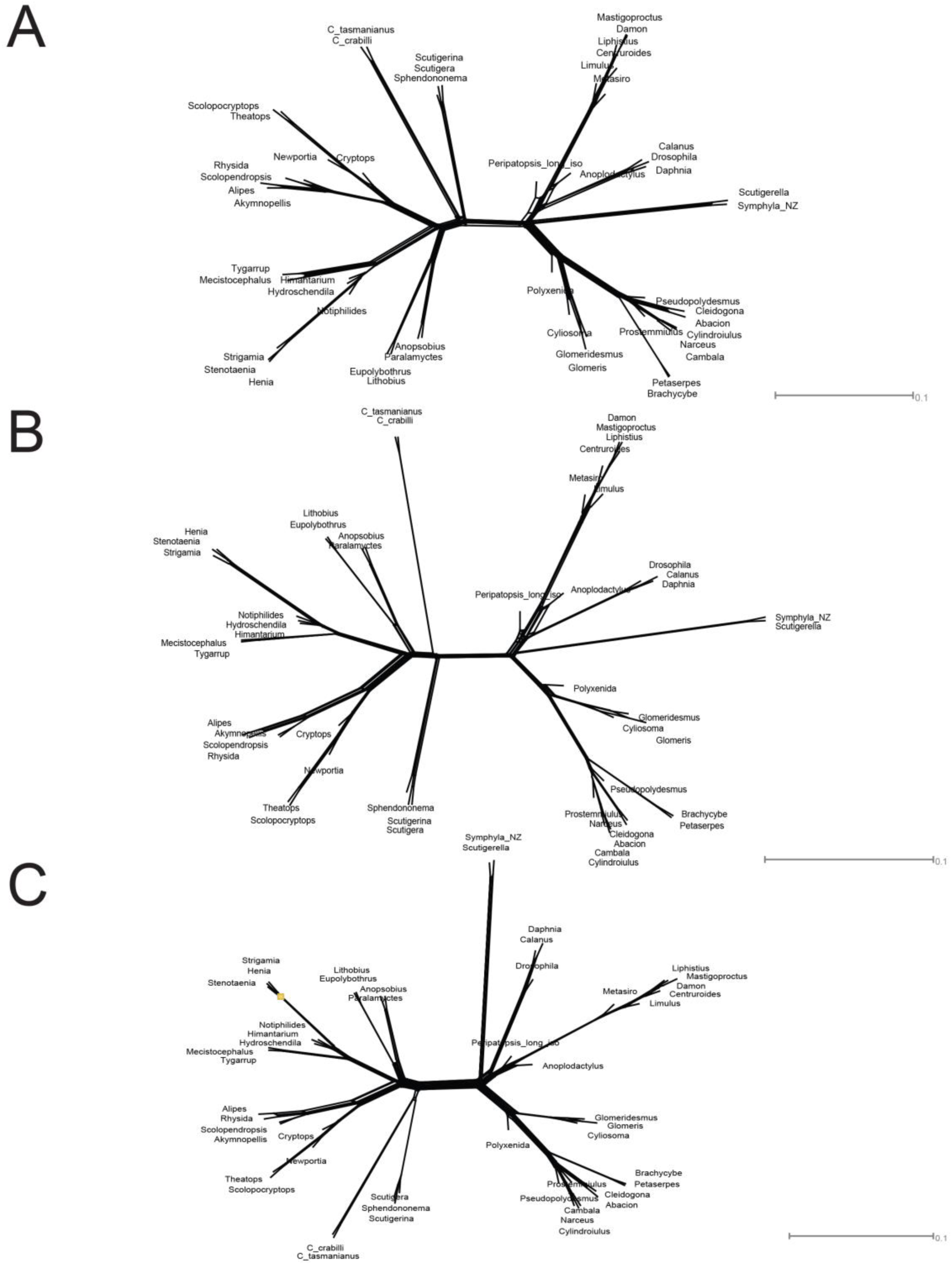
Supernetwork representation of quartets derived from individual ML gene trees, for the genes concatenated in supermatrices I (a), II (b) and III (c). Phylogenetic conflict is represented by reticulations and short branches.

